# splatPop: simulating population scale single-cell RNA sequencing data

**DOI:** 10.1101/2021.06.17.448806

**Authors:** Christina B. Azodi, Luke Zappia, Alicia Oshlack, Davis J. McCarthy

## Abstract

With improving technology and decreasing costs, single-cell RNA sequencing (scRNA-seq) at the population scale has become more viable, opening up the doors to study functional genomics at the single-cell level. This development has lead to a rush to adapt bulk methods and develop new single-cell-specific methods and tools for computational analysis of these studies. Many single-cell methods have been tested, developed, and benchmarked using simulated data. However, current scRNA-seq simulation frameworks do not allow for the simulation of population-scale scRNA-seq data. Here, we present splatPop, a new Splatter model, for flexible, reproducible, and well documented simulation of population-scale scRNA-seq data with known expression quantitative trait loci (eQTL) effects. The splatPop model also allows for the simulation of complex batch effects, cell group effects, and conditional effects between individuals from different cohorts.

## Background

Single-cell RNA-sequencing (scRNA-seq) has enabled the high-throughput quantification of gene expression at the level of the individual cell, making it possible to characterize cells in heterogeneous tissues by their cell-type and cell-state. As gene expression is an intermediate between DNA sequence and traits like response to stimuli and disease status, scRNA-seq can provide insights into the cellular context in which stimuli or diseases have an effect. Today, with decreases in costs and improvements in protocols for multiplexing and demultiplexing samples [1, 2, 3], scRNA-seq is being performed at larger and larger scales, including across population-scale cohorts.

An early focus for many scRNA-seq studies was to identify differentially expressed genes (DEGs) between cell-types or cell-states. Now, with population-scale scRNA-seq, cell-type/state specific DEGs can also be identified between individuals from different cohorts. For example, Lawlor *et al*. [4] identified 638 DEGs across eight different cell-types from the pancreatic tissue of non-diabetes and type 2 diabetes donors. Many of the DEGs were unique to a single cell-type and over half were not discovered when the comparisons were performed without accounting for cell-type, highlighting the importance of single-cell level resolution in deciphering the molecular basis of diseases [4].

Beyond characterizing the cellular context for DEGs, population-scale scRNA-seq data promises to improve our ability to study the regulatory basis for these differences. In recent efforts by the GTEx Consortium to map genetic regulatory effects in human tissues, only 43% of disease-associated genetic variants (i.e. hits from genome wide association studies; GWAS) co-locoalized with expression-associated genetic variants (i.e. expression quantitative trait loci; eQTL) [5]. It was further estimated that, across tissues, only an average of 11% of trait heritability could be explained by GTEX cis-eQTL [6]. One reason for the lack of disease-associated eQTL is that cellular context is critical for genetic regulation and that disease-associated eQTL are missed when eQTL are mapped using data from bulk tissue [7]. Pioneering efforts to use scRNA-seq data for single-cell eQTL (sc-eQTL) mapping have discovered novel cell-type and dynamic-state specific eQTL [8, 9], suggesting population-scale scRNA-seq data could help uncover context-specific regulatory effects.

When scRNA-seq technologies were first becoming available, there was a rush to adapt bulk RNA-seq analysis methods and to develop new single-cell specific analysis methods to address challenges associated with single-cell expression data (e.g., noise, sparsity, high dimensionality). By June 2021, there were over 960 software packages available for scRNA-seq analysis [10]. Many of these new and old methods have been benchmarked to critically assess their performance on a wide variety of data types and conditions, including benchmarks focused on batch-effect correction [11], normalization [12], differential expression [13], and trajectory inference [14]. While the gold-standard for assessing performance of different methods is how well they perform on real datasets, such an assessment can be difficult for scRNA-seq because the ground-truth is typically not known. One solution is to use simulated datasets where the ground truth is known. Splatter, a software package that implements a number of methods to simulate scRNA-seq data (including its own model, splat), has become a popular option for generating realistic simulated scRNA-seq data since its release in 2017 [15]. Splatter is flexible, fast, reproducible, and well maintained and, further, was considered a top-tier model for simulating single-cell RNA-seq data in a recent independent benchmark [16]. However, it does not model genetic effects on expression (i.e., eQTL), precluding its use for population-scale studies. Indeed, we are not aware of any existing scRNA-seq simulation tools that incorporate genetic effects.

Here we present splatPop, an extension of splat for the simulation of population-scale scRNA-seq data with realistic population structure and known eQTL effects. The splatPop model utilizes the flexible framework of Splatter to simulate data with complex experimental designs, including designs with batch effects, multiple cell groups (e.g., cell-types), and individuals with conditional effects (e.g., disease status or treatment effects). These group and conditional effects are simulated by including both DEGs and eQTL effects, making these simulations a powerful tool for assessing downstream single-cell analysis methods. We first present the splatPop model, then demonstrate how splatPop can simulate data with similar properties to simple and complex empirical data sets. Finally, we demonstrate how these simulations can be used to assess single-cell analysis methods. The splatPop model is implemented and available for use in the Splatter package (version 1.16+) available in Bioconductor.

## Results

### The splatPop model

The splatPop framework consists of three steps: (1) estimating parameters from empirical data, (2) simulating population-wide gene means and (3) simulating counts for each cell for each individual, where steps 2 and 3 represent the gamma-Poisson hierarchical modeling approach used in splat.

Three types of splatPop parameters are estimated from empirical data: single-cell, population-scale, and eQTL effects (**Table S1**; **Figure 1, bottom right**). Single-cell parameters (e.g., library size) have previously been described [15], and should be estimated from scRNA-seq data from a homogeneous cell population from one individual. Population-scale parameters (e.g., population-wide expression variance) can be estimated from population-scale bulk RNA-seq or aggregated scRNA-seq data. Finally, eQTL effect size parameters are estimated from real eQTL mapping results. Sensible default parameter values are provided (see **Methods**), however custom values can be estimated from user-provided data or set manually. All parameters are stored in a convenient object that, together with the genotype data, is sufficient to recreate the simulated population.

**Figure 1.**
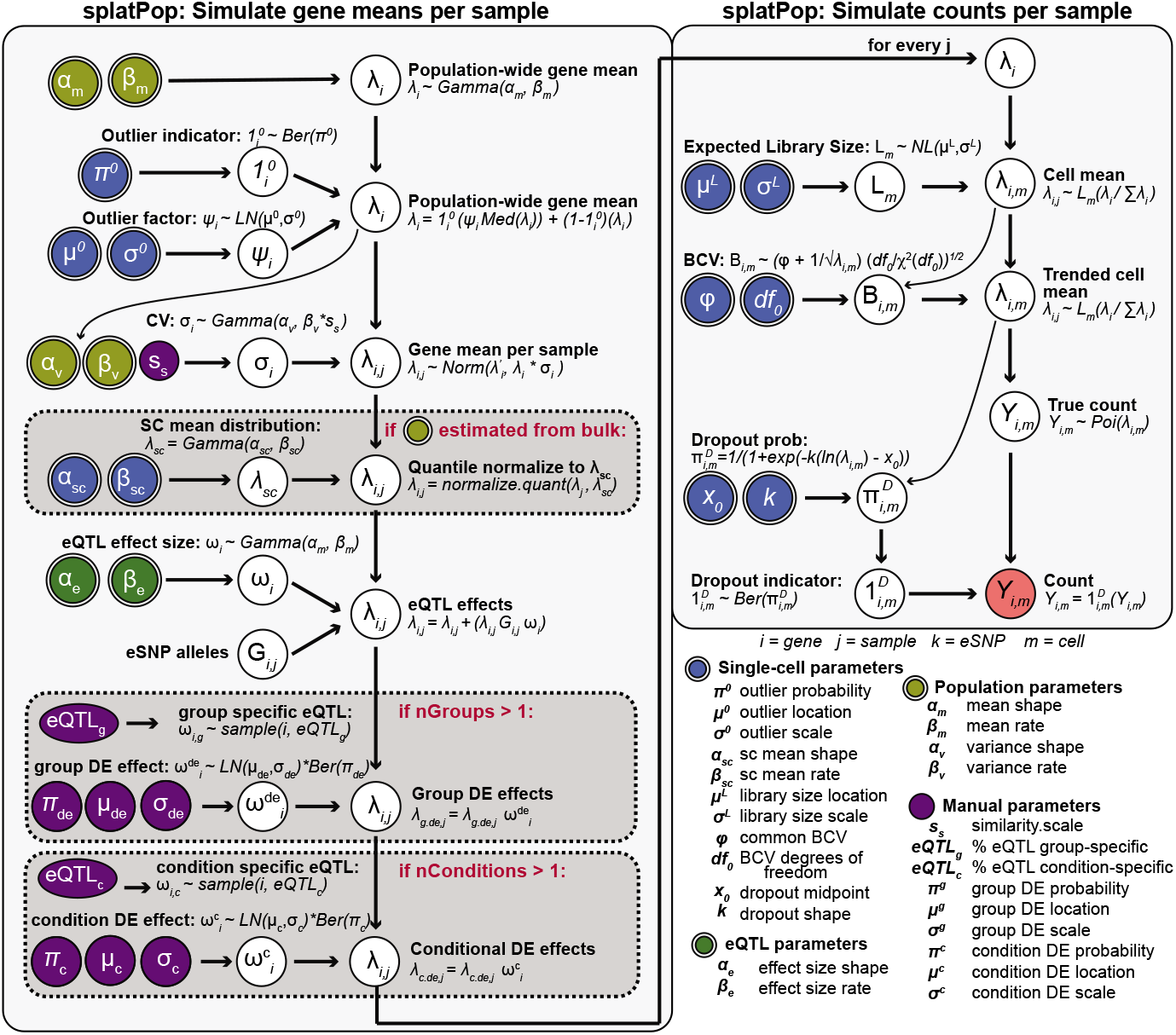
The splatPop model. Input parameters that can be estimated from real data are indicated with double borders and are colored according to type. Steps in gray boxes are optional, with the condition shown in red. (cv: coefficient of variation, BCV: Biological Coefficient of Variation, Ber: Bernoulli, Med: median, LN: log-normal, Poi: Poisson)

To simulate mean expression levels for every gene for every individual (**Figure 1, left panel**), first population-wide means and variance levels are sampled for each gene. To account for the mean-variance trend (**Figure S1**), variance levels are sampled from gamma distributions parameterized from empirical genes in the same gene mean bin, with an option to manually tune the variance between individuals using the *similarity.scale* parameter. Baseline gene means for each individual are then sampled from a normal distribution using the mean and variance assigned to each gene. If population-scale parameters were estimated from bulk data, the baseline means for each individual are quantile normalized to match the desired single-cell distribution. Finally, eQTL and differential expression effects are added to the baseline gene means (see **Methods**). Flexible controls for defining the frequency, size, and context of eQTL and DE effects allow for the simulation of a wide range of complex datasets.

The final step is to simulate realistic expression counts for single-cells for each individual. This task is essentially done using the splat model, where counts are sampled from a Poisson distribution after adjusting means based on the expected library size and BCV (**Figure 1, right panel**). The splatPop model also uses the batch effect function from splat to simulate population-scale data from multiplexed experimental designs with technical replicates (i.e. where all or some individuals are simulated in more than one batch).

### Simple splatPop simulations

Three empirical datasets are used as reference data throughout our study: induced Pluripotent Stem Cells (iPSCs) captured on the SmartSeq2 platform (ss2-iPSC; [9]), floor plate progenitor and dopaminergic neuron cells from a panel of iPSCs differentiating toward a midbrain neural fate captured on the 10x platform (10x-Neuro; [17]), and fibroblasts from models of idiopathic pulmonary fibrosis (IPF) captured on the 10x platform (10x-IPF; [18]; see **Methods**). For each reference, single-cell parameters were estimated using the individual with the most cells (ss2-iPSC: joxm n=383, 10x-Neuro: wihj n=2268, 10x-IPF: 947170 n=399) and population parameters were estimated from mean aggregated counts, excluding individuals with less than 100 cells and individuals from the disease cohort for 10x-IPF.

First, using the ss2-iPSC data as a reference, we simulated scRNA-seq counts for genes on chromosome 22 (504 genes) for six individuals from the 1000 Genomes Project. This example simulation took less than 30 seconds to generate, but splatPop can be efficiently scaled up. For example, simulating 1,000 genes for 1,000 individuals takes just over five minutes (**Table S2**. The *similarity.scale* parameter and the percentage of genes assigned with eQTL effects were manually adjusted to match data from six individuals from the reference that were sequenced in the same batch.

Since Zappia *et al*. and the independent benchmark by Cao *et al*. demonstrate splat’s performance compared to other scRNA-seq simulation models in terms of various metrics at the individual-level [15, 16], here we focus on population-level characteristics between splatPop simulations and empirical data. For example, visualizing the global relationship between cells in a lower-dimensional space (**Figure 2a**), we see that both the empirical and simulated cells loosely cluster by individual, but with significant overlap. We can quantify the degree of clustering by individual by calculating the silhouette width of each cell, a measure of its similarity to other cells from the same individual compared to cells from its nearest neighbor individual. While simulations from a parametric model cannot perfectly replicate empirical data, we find a similar distribution of silhouette widths across simulated compared to empirical cells (**Figure 2b, left**). Separating the silhouette widths by individual, we see that cells from some individuals cluster more distinctly than others (**Figure 2b, right**). Note that the simulated individuals are not the same genotypes as the reference individuals, thus we do not expect the same inter-individual relationship pattern to be observed between simulated and empirical cells. However, such relationship patterns could be achieved by providing splatPop with the genotype data for the population of interest.

**Figure 2.**
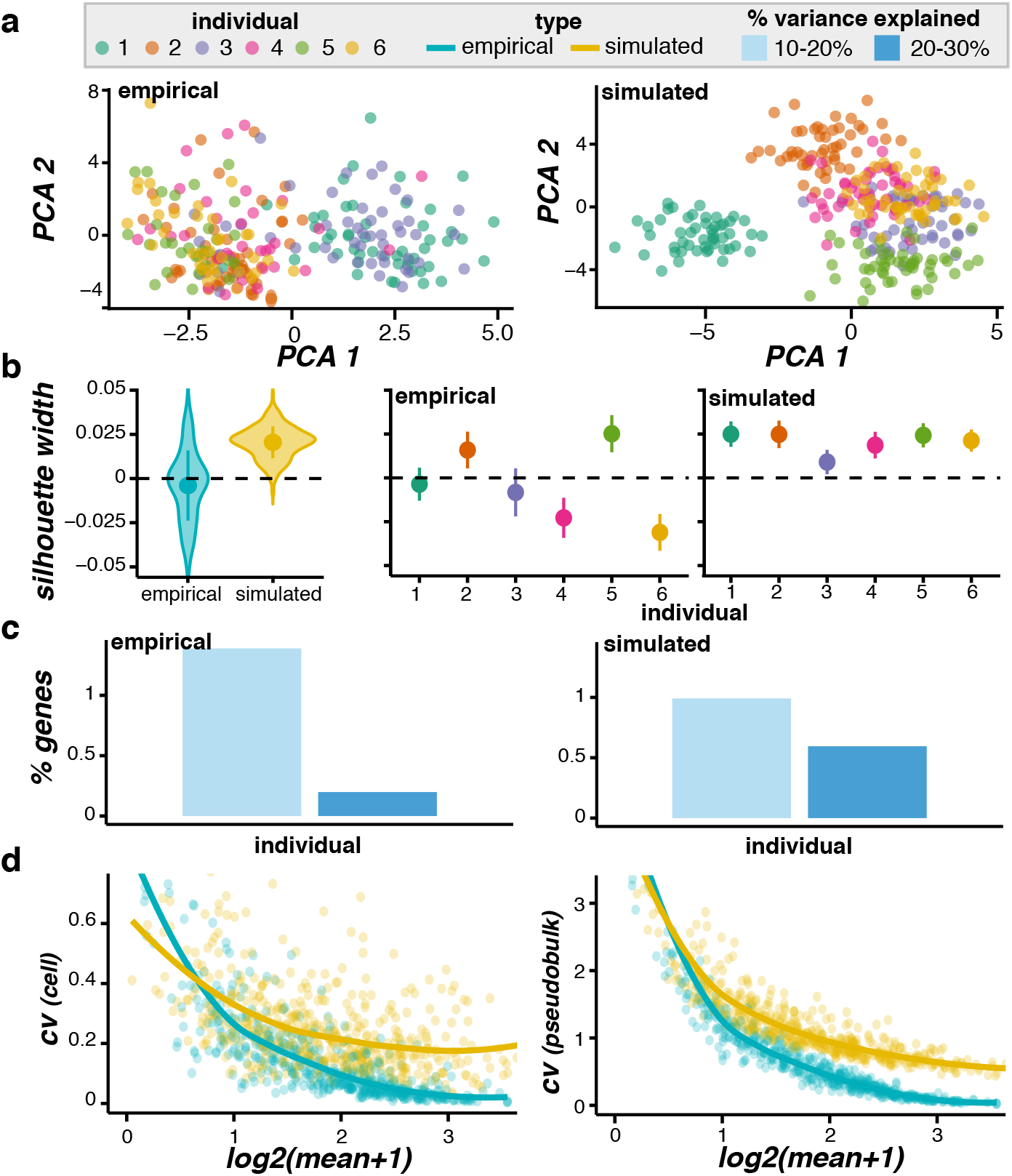
Simulated data compared to ss2-iPSC empirical scRNA-seq data. (**a**) PCA plots of cells colored by individual (max 50 cells shown per individual). (**b**) The distribution of cell silhouette widths using the individual as the cluster. The distributions are shown for cells grouped by type (left) and by type and individual (right), with the point and whisker showing the mean and standard deviation. (**c**) The percentage of genes (y-axis) with a given percentage of variance explained by individual. (**d**) The mean-variance relationship, with the average gene mean (x-axis) and the gene coefficient of variation (cv; y-axis) calculated across (left) all cells or (right) pseudo-bulk data aggregated by individual. Empirical: n = average 52 cells per individual; Simulated: n = 52 cells per individual.

In addition to these cell-level comparisons, we can also compare our simulated data with the reference data in terms of gene-level characteristics. By calculating the percentage of variance in gene expression across cells explained by an experimental factor, such as individual, we see that for both the reference and simulated data, nearly 2% of genes have between 10-30% of their variance explained by individual (**Figure 2c**). Another key aspect of scRNA-seq data is the mean-variance relationship. Here we compare the mean-variance relationship for each gene across all cells and across pseudo-bulk data aggregated by individual (see **Methods**; **Figure 2d, left and right, respectively**). We also demonstrate that splatPop can simulate a simple data set with the cell-level and gene-level properties of data generated on the 10x sequencing platform (**Figure S2**). Together, these comparisons demonstrate that splatPop can approximate the cell and gene level characteristics of single-cell RNA-sequencing data.

### Complex simulations

The simulations above demonstrate how splatPop can provide population scale scRNA-seq data for a homogeneous cell population where the expression differences between individuals are only due to random sampling and genetic effects. Next we demonstrate how splatPop can be used to simulate data with more complex effects (**Figure 3)**.

**Figure 3.**
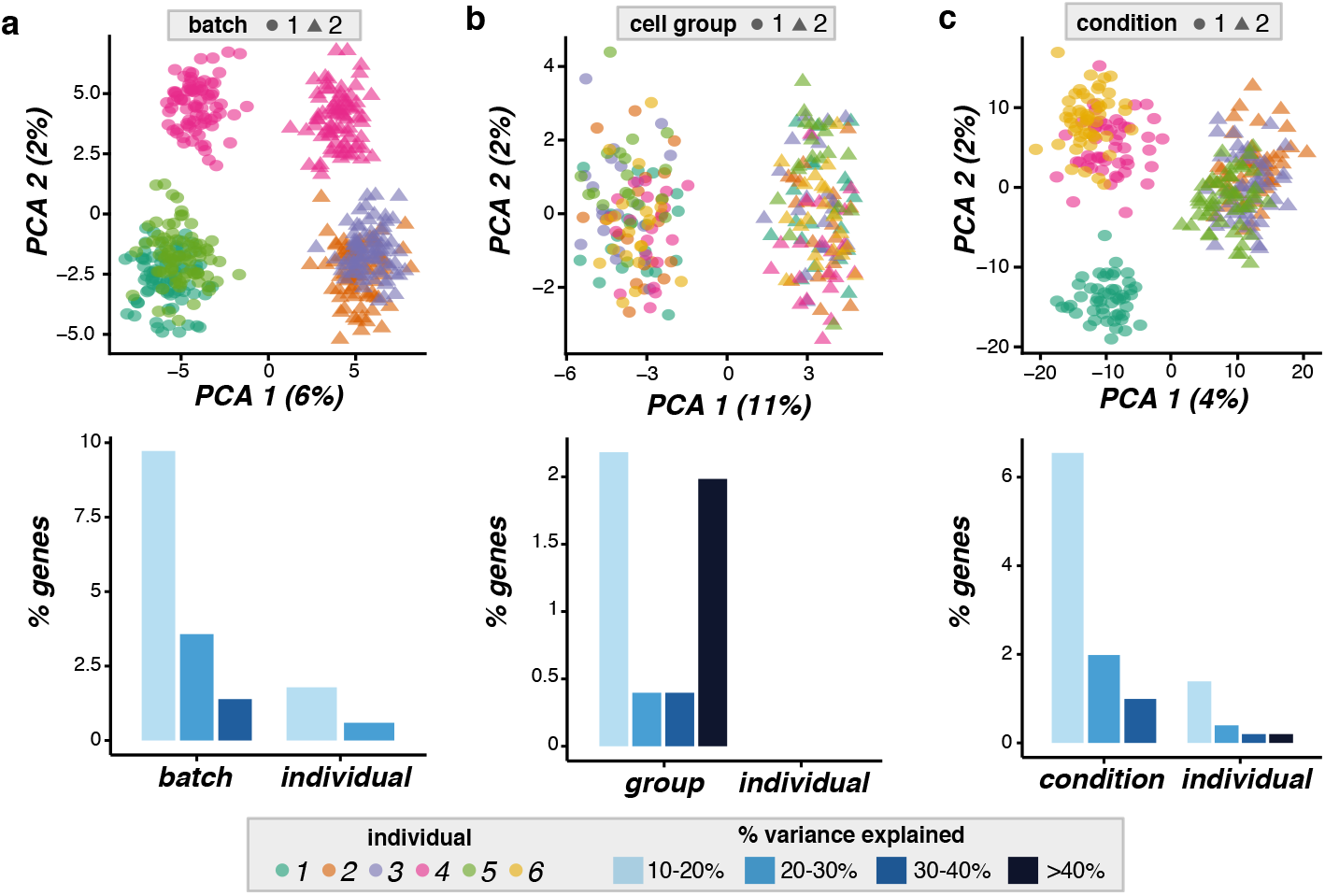
Examples of complex splatPop simulations. A principal component analysis (PCA; top) plot of the simulations with each individual designated by color and each complex effect denoted by shape and the distribution of the degree of variance in gene expression across cells explained by the experimental factor (bottom) for simulations with **(a)** batch effects using ss2-iPSC as a reference, **(b)** cell group effects using 10x-Neuro as a reference, and **(c)** conditional effects using 10x-IPF as a reference. Note, no variance in gene expression is explained by individual using the 10x-Neuro reference.

#### Simulating batch effects

Unwanted sources of variation due to technical differences during sample collection, processing, and sequencing are known as batch effects. Batch effects are especially important to consider for population-scale scRNA-seq simulations because these data are often generated by combining multiple individuals together by pooling or multiplexing. The splat model simulates batch effects by sampling a multiplicative effect for each gene for each batch and applying it to all cells in the batch. In splatPop, we use this same approach, but expand the function to allow the user to design complex multiplexing schemes.

We demonstrate this function by simulating five individuals in two batches using the ss2-iPSC data as a reference. Batch effect sizes are sampled from a log-normal distribution for each gene. Here we adjust the location and scale parameters to simulate a data set where the batch effects are larger than the individual effects. Because we specified for three individuals to be included in each batch, splatPop randomly selected one individual (sample 4) to be simulated in both batches (**Figure 3a**). As with the simple simulations, we can also tune the *similarity.scale* parameter, percentage of genes assigned as eQTL, and the location and scale parameter for the batch effects to simulate data that closely resembles empirical data. For example, **Figure S3** shows a full comparison between ss2-iPSC reference data from 10 individuals sequenced across 3 batches. In this example, the batch effect parameters were set individually for each batch so that one batch (batch 3) had larger batch effects than the other batches, highlighting the flexibility of splatPop.

#### Simulating cell groups

Multiple cell groups can be simulated to mimic heterogeneous cell populations (i.e. cell types) for each individual or to mimic treated verses untreated cells from the same individual. The splat model simulates cell groups by assigning group-specific multiplicative DE factors, sampled from a log-normal distribution, to a subset of genes. In splatPop, in addition to the DE factors, we also randomly designate a proportion of eQTL effects as cell group specific. In this way, different cell groups are defined by genetic and non-genetic DE. The proportions of genes with group-specific eQTL and DE and the level of DE can be set for each group, allowing for the simulation of highly complex cell populations.

We demonstrate this function by simulating two cell groups for six individuals using the 10x-Neuro data as a reference. Because the 10x-Neuro data had weak individual effects (i.e. cells from different individuals all clustered together), the group effects dominate this simulation (**Figure 3b**). We can tune the splatPop parameters to simulate a dataset with cell group effects that closely resembles empirical data (**Figure S4**).

#### Simulating conditional effects

Finally, a desirable feature for a population-scale scRNA-seq simulation framework is the ability to simulate differences between individuals that are due to different treatments or conditions (e.g., disease status). The splatPop model allows the user to define the number of conditional groups and the proportional of individuals assigned to each group, and then applies condition-specific DE and eQTL effects. This approach is similar to how cell groups are simulated, but effects are applied to all cells for all individuals in the condition group. For example, group effects can be used to simulate both treated and untreated cells for all individuals in the population, while conditional effects can be used to simulate cells for treated and untreated individuals. Again, the proportion of condition-specific-eQTL and the proportion of genes with condition-specific DE and the level of DE can be specified separately for each conditional group.

We demonstrate this function by simulating six individuals, three in each conditional group, using the 10x-IPF data as a reference. We adjust the location and scale parameters defining the DE effect sizes and proportion of condition-specific eQTL to simulate a data set where the conditional effects are larger than the individual effects (**Figure 3c**). Again, we can tune these parameters to simulate datasets with conditional effects that closely resemble empirical data (**Figure S5**).

### Using splatPop to evaluate downstream analysis tools

The simulations from splatPop can be used to evaluate new and existing scRNA-seq analysis methods [19]. They can also be used to assess power to detect known effects across experimental variables, such as number of cells sequenced per individual. Here we demonstrate splatPop’s capabilities with example evaluations of approaches to differential expression analysis and eQTL mapping.

For DE analysis, we simulated 500 cells for each of 12 individuals divided into two conditional groups (6 per group) using 10x-IPF as a reference. Conditional DE effects were assigned randomly to 40% of genes, with scaling factors sampled from a log normal distribution (location = 0.2, scale = 0.2). We then tested for DE genes between the two conditional groups by applying Wilcoxon rank sum tests to pseudo-bulked counts (i.e., sum aggregation of counts across cells for each individual) calculated from down-sampled subsets of the cells ranging from 10 to 500 cells per individual. Because we know which genes were simulated with conditional DE effects, we can calculate performance metrics such as true positive rate (TPR) and false discovery rate (FDR) (**Figure 4a**). We also took a closer look at the properties of simulated genes that were correctly identified as DE (true positives; TP) compared to genes that were missed (false negative; FN) or incorrectly identified as DE (false positives; FP) using 80 cells per individual. TP genes tended to have higher mean expression and tended to have larger DE effect sizes (**Figure 4b**).

**Figure 4.**
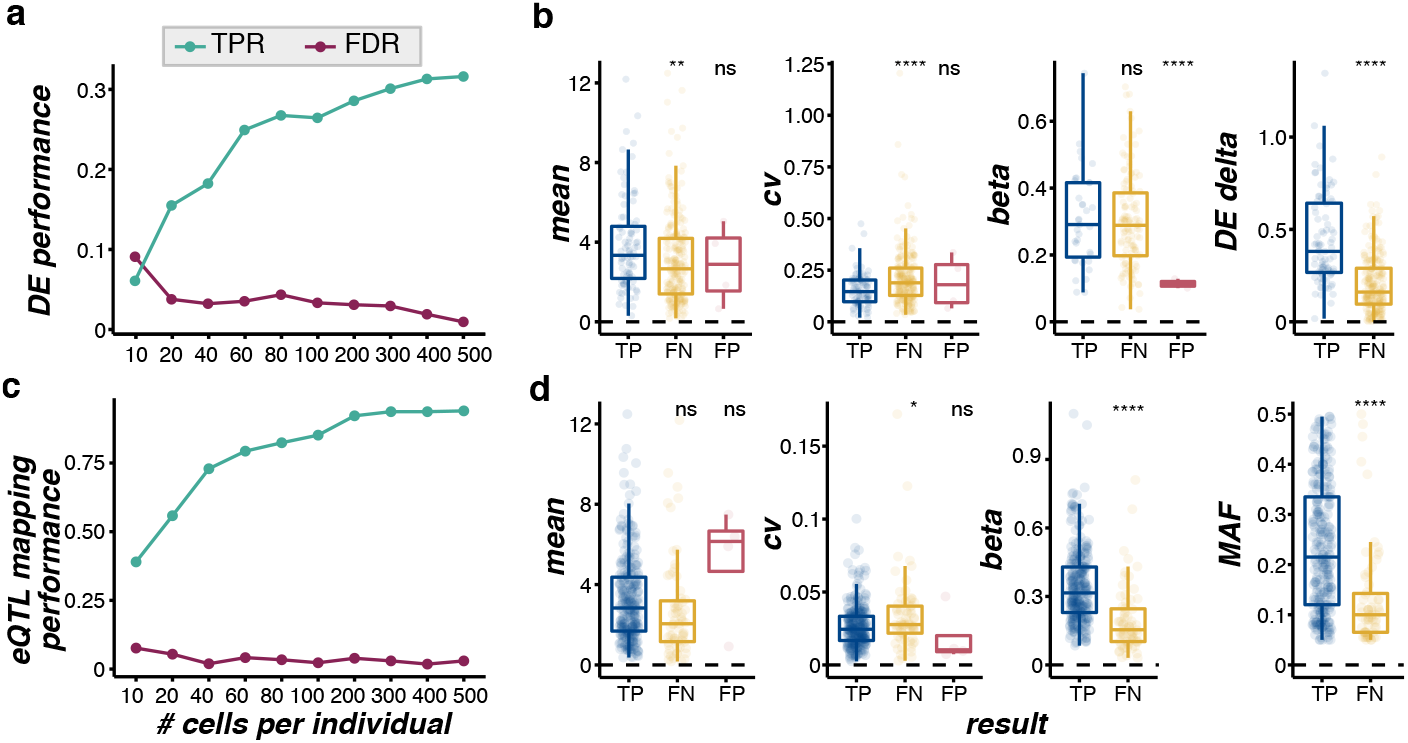
Example applications of splatPop simulations. Pseudo-bulk differential expression (DE) analysis: (**a**) The true positive rate (TPR: TP/(TP+FN)) and false discovery rate (FDR: FP/(TP+FP)) of DE genes (Wilcoxon rank sum test q.value < 0.1) between two conditional groups across a range of number of simulated cells per individual using 10x-IPF as a reference. (**b**) The simulated gene mean and coefficient of variation (cv), eQTL effect size (beta, if applicable to that gene), and DE effect size (DE delta) for TP, FN, and FP DE genes using 80 cells per individual. (**c**) The performance of eQTL mapping (q-value < 0.05) across a range of number of simulated cells per individual. (**d**) The simulated gene mean and coefficient of variation, eQTL effect size (beta), and eSNP minor allele frequency (MAF) for TP, FN, and FP eGenes using 80 cells per individual. Statistical significance is reported for t-tests testing for difference between TP and FN or FP categories (ns: p > 0.05, *: p <= 0.05, **: p <=, 0.01, ***: p <= 0.001, ****: p <= 0.0001)

For eQTL analysis, we simulated 500 cells for each of 100 individuals in 10 batches with 10 individuals per batch using ss2-iPSC as a reference. We assigned eQTL effects to 70% of simulated genes. We used the optimized single-cell eQTL mapping workflow defined by Cuomo, Alvari, Azodi, *et al*. [19]) to map eQTL (see **Methods**) and assessed performance as the number of cells per individual used increased from 10 to 500 (**Figure 4c**). Again, we took a closer look at the properties of TP, FN, and FP eQTL associations using 80 cells per individual. eQTL tended to be missed for genes with higher coefficient of variation and for eQTL with small effect sizes and when the variant assigned to the gene had a low minor allele frequency (**Figure 4d**). While a thorough benchmarking of approaches for DE analysis or eQTL mapping is beyond the scope of this paper, these examples demonstrate the utility of splatPop for assessing population-scale single-cell analysis methods and experimental design considerations.

## Discussion and conclusions

The expansion of single-cell RNA sequencing to population-scale cohorts has powerful implications for studies of functional genomics. New tools and methods need to be rapidly developed and rigorously tested to ensure researchers are able to make important new discoveries from these data. Independent simulation frameworks are a critical resource to this end.

Here we present splatPop, a flexible framework for simulating population-scale single-cell RNA-sequencing data using the splat model. splatPop is implemented and available in the Splatter R package from Bioconductor, under a GPL-3 license. Because the splatPop model simulates genetic effects on gene expression (i.e. eQTL), realistic population structure can be achieved by providing real genotype data to the model. In addition to being able to simulate complex batch effects and cell group effects, splatPop allows for the simulation of conditional (e.g., treatment or disease status) effects for different cohorts of individuals. The modular framework of splatPop enables these functions to be combined to generate synthetic data with complex experimental designs. For example, splatPop could be used to simulate data for multiple cell-types for individuals from healthy, diseased, and disease-treated individuals, sequenced using a multiplex design with technical replicates.

We demonstrate that by estimating splatPop parameters from real data and manually adjusting control parameters, synthetic data can be generated that has properties resembling a wide range of real data sets. As a parametric simulation framework, splatPop cannot perfectly reproduce all aspects of empirical scRNA-sequencing data, but it has the benefits of flexibility, speed, and parameter interpretability. The simulation functions available in splatPop are well documented and reproducible. Code used to generate all simulations shown here and to generate the plots comparing the simulated to reference data are also available. Further, as improvements are made to the splat model, they will be adopted by splatPop.

## Methods

### Reference data

Empirical scRNA-seq data sets used as references are all previously described and available for download as processed, post-quality control counts per cell per gene (ss2-iPSC [9]: https://zenodo.org/record/3625024#.YC3NyZMzZTY; 10x-Neuro [17]: https://zenodo.org/record/4333872#.YFAM-UgzZTa; and [18]: https://www.ebi.ac.uk/gxa/sc/experiments/E-HCAD-14/downloads).

Additional cell-level and gene-level filtering was performed as follows. For ss2-iPSC, only cells sequenced at day 0 were included (i.e. iPSCs). For 10x-Neuro, only cells from day 30 that were annotated as floor plate progenitors or dopaminergic neurons and were sequenced in the largest pool (pool 7) were included. Gene filtering was also done to remove genes with zero variance or a mean across cells equal to zero. For 10x-IPF, only fibroblast annotated cells from control and bleomycin 1.7 mg/kg fibrosis-induced mice were included. Annotation was done using SingleR v1.4.1 with the MouseRNAseqData cell type reference from CellDex v1.2.0 [20]. splatPop was estimating very large variance parameters (biological coefficient of variation common dispersion greater than 10) from 10x-IPF. This behavior was driven by the unusually deep sequencing of the libraries for this study, resulting in counts per gene values across cells that often reached into the hundreds while also exhibiting a considerable number of observed zeros. Filtering out genes with zero variance and zero mean as was done for the 10x-Neuro reference was not sufficient to remove this effect. Therefore, we performed additional filtering whereby cells were removed if they had zero counts for over 95% of genes and genes were removed if they had zero counts for over 60% of the remaining cells.

For each reference, the genes included were randomly down-sampled to n = 504 (after other gene filtering steps for the two 10x references) to match the number of genes being simulated (i.e., number of genes on chromosome 22). Single-cell parameters were estimated using the individual with the most cells (ss2-iPSC: joxm n=383, 10x-Neuro: wihj n=2268, 947170: n=399) and population parameters were estimated from mean aggregated counts (i.e., taking the mean of the count values across the cells for each gene from each individual), excluding individuals with less than 100 cells and individuals from the disease cohort for 10x-IPF. The eQTL effect size parameters were estimated from eQTL mapping beta estimates for the top eQTL hit for each gene (for SNPs with MAF greater than 10%) from GTEx v7 using thyroid tissue [21]. These are the default splatPop eQTL effect size parameters.

Genotype data from the 1000 Genomes project for chromosome 22 was used for the simulations (hg19, phase3 [22]). The genotype data was filtered to include biallelic SNPs with no missing data, with minor allele frequency greater than 5%, Hardy-Weinberg equilibrium exact test p-value greater than 0.00001, and to remove SNPs in high linkage disequilibrium (independent pairwise r2 less than 0.75, 1600 kb window).

### splatPop simulation framework

The splatPop framework (**Figure 1)** consists of (i) estimating key parameters from empirical data, (ii) simulating gene means for the population, and (iii) simulating single-cell counts for the population.

#### Step 1. Parameter estimation

The estimation of single-cell parameters is described in detail in the original Splatter manuscript [15]. Population parameters are estimated from population scale bulk RNA-seq or aggregated scRNA-seq data, provided as a matrix of expression levels for each gene (rows) for each individual (column). The parameters that control population-wide mean expression of each gene (*α_m_* and *β_m_*) are estimated by fitting a gamma distribution to the gene means across the population. Lowly-expressed genes (expression less than 0.1 in greater than 50% of individuals) are excluded. To account for the mean-variance relationship (**Figure S1**), parameters that control expression variance across the population (*α_υ_* and *β_υ_*) are estimated by fitting a gamma distribution to the coefficient of variation (cv) from genes in each expression bin (default n.bins=10). The parameters that control the eQTL effect sizes (*α_e_* and *β_e_*) are estimated by fitting a gamma distribution to effect sizes from an empirical eQTL mapping study (bulk or single-cell).

The default values for the population and eQTL parameters were estimated from bulk data from GTEx (v7, thyroid tissue) [21]. The effect sizes from the top eSNP for each gene were used, after removing top eSNPs with a MAF less than 0.05.

#### Step 2. Simulating gene means for individuals

Population-wide expression means are modeled as *λ_i_* ~ *gamma*(*α_m_, β_m_*) for genes *i* = {*i*_1_, …, *i_n.genes_*}. If specified by the single-cell parameters, expression outlier effects are added to the sampled values as described in Zappia *et al*. [15]. A variance is sampled for each gene as *σ_i_* ~ *gamma*(*α_υ_, β_υ_*), where *α_υ_* and *β_υ_* are specific to the gene mean bin. The variance can be manually adjusted with the *similarity.scale s_υ_* parameter, which gets multiplied with the *β_υ_* parameter. The baseline gene means per individual are then modeled as *λ_i,j_* ~ *N* (*λ_i_, λ_i_* * *σ_i_*) for every individual *j* = {*j*_i_, …, *j_individuals_*}. If the population-scale parameters were estimated from bulk RNA-seq data, for each individual *j*, the sampled means *λ_i_* are quantile normalized to match the distribution estimated from the single-cell data (i.e., *gamma*(*α_sc_, β_sc_*)).

The simulation of eQTL effects requires genotype information as input. Providing real genotype data or genotype information simulated using tools like HAPGEN2 [23] will ensure the simulated data reflects realistic population structure. Various control parameters can also be adjusted, including how many or what proportion of genes are assigned eQTL effects (i.e. eGenes), the minimum and maximum minor allele frequency (MAF) of the SNP assigned to each eGene (i.e. eSNP), and the maximum distance between eGenes and eSNPs. For each eGene, an effect size is sampled as *ω_i_* ~ *gamma*(*α_e_, β_e_*). The eQTL effects are incorporated into the baseline means as in [24] using the equation:

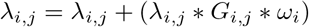

where *G_i,j_* is the minor allele dosage of the eSNP assigned to eGene *i*, coded as 0, 1, or 2, for individual *j*. To account for cell-group differences, a portion of eQTL effects (specified by *eqtl.group.specific*) are applied to the baseline means for the cells simulated as belonging to a specific cell-group. To account for cohort differences, a portion of eQTL effects (specified by *eqtl.condition.specific*) are only applied to the baseline means for individuals assigned to a specific conditional cohort.

DE effects between cell-groups or between conditional cohorts are simulated as in [15]. Briefly, DE scaling factors are sampled as 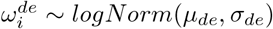 for the genes assigned DE effects (proportion specified by *de.prob* and *cde.prob*), where *μ_de_* and *σ_de_* can be adjusted separately for cell-groups and cohorts to change the relative impact of these effects. A portion of DE effects can be randomly assigned as negative effects (specified by *de.downProb* and *cde.downProb*).

#### Step 3. Simulating gene means for individuals

Single-cell level expression counts are simulated using the splat model [15], with minor modifications to account for splatPop having sampled gene means and incorporated expression outliers in step 2. Batch effects are also incorporated into the simulations during this step, where multiplicative factors are added to genes for cells from the same batch. In splatPop, the batch effect function from splat is expanded to allow for the simulation of complex multiplexed experimental designs, where individuals are pooled where the multiplicative factors are applied to all cells from all individuals in that batch. The user can specify the number of batches (length of list provided to *batchCells*) and the number of individuals-per-batch (*batch.size*). By adjusting these parameters, splatPop can simulate populations where there are no batches, where all individuals are present in multiple batches, or where a subset of individuals are present in multiple batches as technical replicates.

### Comparing simulated and empirical data

Principal component analysis was performed and plotted using functions from scater v1.18.6 [25]. Silhouette widths for each cell were calculated using the silhouette function from R package cluster v2.1.0 [26], using the Euclidean distance between normalized log counts. For each cell, the silhouette width was calculated using the individual as the cluster, and when applicable with the batch, cell group, or conditional group as the clustering factor(s). The percentage of gene expression variance explained and aggregated count values were also calculated using functions from scater.

### Differential expression and eQTL mapping analyses

For DE analysis, scRNA-seq data was simulated using 10x-IPF as a reference for 12 individuals in two conditional groups (six per group), with 500 cells per individual. Pseudo-bulk counts were generated by sum aggregating N single-cell counts across individuals and applying library size normalization, for N ranging from 10 to 500. The Wilcoxon rank sum test was applied and p-values were corrected for multiple testing using estimated Benjamini-Hochberg FDR [27] with q-value less than 0.1 as a threshold for DE significance.

The eQTL mapping was performed following the optimized single-cell eQTL mapping workflow defined by Cuomo, Alvari, Azodi, *et al*. [19]. Briefly, single-cell level normalization is performed on the simulated counts using scran v1.20 [28], then counts are mean aggregated by individual and quantile normalized to a standard normal distribution. All SNPs within 100kb up- and down-stream of the gene are tested using a linear mixed model fit using LIMIX [29]. In addition to the SNP fixed effect, the LMM includes the top 15 principal components from a PCA on the expression data as fixed effects to account for unwanted variation and the kinship (realized relationship matrix calculated using plink v1.90 [30]) as a random effect to account for population structure. Gene-level p-values are controlled for multiple testing using the empirical null distribution from 100 permutations of the genotype data. Then across-gene multiple testing correction is performed on the top SNP per gene using the Storey Q value procedure and the p-value corresponding to the q < 0.05 threshold is determined. All eQTL associations with p-value below this level are then considered significant.

To evaluate DE and eQTL mapping we calculated the number of true positives (TP), true negatives (TN), false positives (FP), and false negatives (FN) and summarize these into true positive rate (TPR: TP/(TP+FN)) and false discovery rate (FDR: FP/(TP+FP)).

## Availability

splatPop is available in Splatter v1.16+ on Bioconductor at https://bioconductor.org/packages/release/bioc/html/splatter.html and on GitHub at https://github.com/Oshlack/splatter. All of the code used to generate the simulations and analyses in this manuscript are available at https://biocellgen-public.svi.edu.au/KEJP_2020_splatPop/public/index.html.

## Competing interests

The authors declare that they have no competing interests.

## Author’s contributions

D.J.M and C.B.A. conceived the study. C.B.A developed the splatPop software and conducted the study. L.Z. and A.O. developed the original Splatter software and contributed to splatPop software development. D.J.M. supervised all aspects of the project. All authors contributed to, read, and approved the final manuscript.

## Acknowledgements

We would like to thank Marc Jan Bonder, Anna S.E. Cuomo, and Jeffrey Pullin for useful discussions regarding simulating population scale-scRNA-seq data and for their help with software testing.

## Funding

C.B.A. is supported by funding from the Baker Foundation. A.O is supported by the National Health and Medical Research Council (NHMRC) of Australia through an Ideas grant (GNT1187748) and an Investigator Grant (GNT1196256). D.J.M. is supported by the NHMRC through an Early Career Fellowship (GNT1112681), a Project Grant (GNT1162829), and an Investigator Grant (GNT1195595), by the Baker Foundation, and by Paul Holyoake and Marg Downey through a gift to St Vincent’s Institute of Medical Research.

## Notes

https://biocellgen-public.svi.edu.au/KEJP_2020_splatPop/public/index.html

### Competing Interest Statement

The authors have declared no competing interest.

## References

1. McCarthy, D.J., HipSci Consortium, Rostom, R., Huang, Y., Kunz, D.J., Danecek, P., Bonder, M.J., Hagai, T., Lyu, R., Wang, W., Gaffney, D.J., Simons, B.D., Stegle, O., Teichmann, S.A.: Cardelino: computational integration of somatic clonal substructure and single-cell transcriptomes (2020)

2. Huang, Y., McCarthy, D.J., Stegle, O.: Vireo: Bayesian demultiplexing of pooled single-cell RNA-seq data without genotype reference. Genome Biol. 20(1), 273 (2019)

3. Kang, H.M., Subramaniam, M., Targ, S., Nguyen, M., Maliskova, L., McCarthy, E., Wan, E., Wong, S., Byrnes, L., Lanata, C.M., Gate, R.E., Mostafavi, S., Marson, A., Zaitlen, N., Criswell, L.A., Ye, C.J.: Multiplexed droplet single-cell RNA-sequencing using natural genetic variation. Nat. Biotechnol. 36(1), 89–94 (2018)

4. Lawlor, N., George, J., Bolisetty, M., Kursawe, R., Sun, L., Sivakamasundari, V., Kycia, I., Robson, P., Stitzel, M.L.: Single-cell transcriptomes identify human islet cell signatures and reveal cell-type-specific expression changes in type 2 diabetes. Genome Res. 27(2), 208–222 (2017)

5. GTEx Consortium: The GTEx consortium atlas of genetic regulatory effects across human tissues. Science 369(6509), 1318–1330 (2020)

6. Yao, D.W., O’Connor, L.J., Price, A.L., Gusev, A.: Quantifying genetic effects on disease mediated by assayed gene expression levels. Nat. Genet. 52(6), 626–633 (2020)

7. Umans, B.D., Battle, A., Gilad, Y.: Where are the Disease-Associated eQTLs? Trends Genet. 37(2), 109–124 (2021)

8. van der Wijst, M.G.P., Brugge, H., de Vries, D.H., Deelen, P., Swertz, M.A., LifeLines Cohort Study, BIOS Consortium, Franke, L.: Single-cell RNA sequencing identifies celltype-specific cis-eQTLs and co-expression QTLs. Nat. Genet. 50(4), 493–497 (2018)

9. Cuomo, A.S.E., Seaton, D.D., McCarthy, D.J., Martinez, I., Bonder, M.J., Garcia-Bernardo, J., Amatya, S., Madrigal, P., Isaacson, A., Buettner, F., Knights, A., Natarajan, K.N., HipSci Consortium, Vallier, L., Marioni, J.C., Chhatriwala, M., Stegle, O.: Single-cell RNA-sequencing of differentiating iPS cells reveals dynamic genetic effects on gene expression. Nat. Commun. 11(1), 810 (2020)

10. Zappia, L., Phipson, B., Oshlack, A.: Exploring the single-cell RNA-seq analysis landscape with the scRNA-tools database. PLoS Comput. Biol. 14(6), 1006245 (2018)

11. Tran, H.T.N., Ang, K.S., Chevrier, M., Zhang, X., Lee, N.Y.S., Goh, M., Chen, J.: A benchmark of batch-effect correction methods for single-cell RNA sequencing data. Genome Biol. 21(1), 12 (2020)

12. Cole, M.B., Risso, D., Wagner, A., DeTomaso, D., Ngai, J., Purdom, E., Dudoit, S., Yosef, N.: Performance assessment and selection of normalization procedures for Single-Cell RNA-Seq. Cell Syst 8(4), 315–3288 (2019)

13. Soneson, C., Robinson, M.D.: Bias, robustness and scalability in single-cell differential expression analysis. Nat. Methods 15(4), 255–261 (2018)

14. Saelens, W., Cannoodt, R., Todorov, H., Saeys, Y.: A comparison of single-cell trajectory inference methods. Nat. Biotechnol. 37(5), 547–554 (2019)

15. Zappia, L., Phipson, B., Oshlack, A.: Splatter: simulation of single-cell RNA sequencing data. Genome Biol. 18(1), 174 (2017)

16. Cao, Y., Yang, P., Yang, J.Y.H.: A benchmark study of simulation methods for single-cell RNA sequencing data (2021)

17. Jerber, J., Seaton, D.D., Cuomo, A.S.E., Kumasaka, N., Haldane, J., Steer, J., Patel, M., Pearce, D., Andersson, M., Bonder, M.J., Mountjoy, E., Ghoussaini, M., Lancaster, M.A., Marioni, J.C., Merkle, F.T., Gaffney, D.J., Stegle, O., HipSci Consortium: Population-scale single-cell RNA-seq profiling across dopaminergic neuron differentiation. Nat. Genet. (2021)

18. Peyser, R., MacDonnell, S., Gao, Y., Cheng, L., Kim, Y., Kaplan, T., Ruan, Q., Wei, Y., Ni, M., Adler, C., Zhang, W., Devalaraja-Narashimha, K., Grindley, J., Halasz, G., Morton, L.: Defining the activated fibroblast population in lung fibrosis using Single-Cell sequencing. Am. J. Respir. Cell Mol. Biol. 61(1), 74–85 (2019)

19. Cuomo, A.S.E., Alvari, G., Azodi, C.B., single-cell eQTLGen consortium, McCarthy, D.J., Bonder, M.J.: Optimising expression quantitative trait locus mapping workflows for single-cell studies (2021)

20. Aran, D., Looney, A.P., Liu, L., Wu, E., Fong, V., Hsu, A., Chak, S., Naikawadi, R.P., Wolters, P.J., Abate, A.R., Butte, A.J., Bhattacharya, M.: Reference-based analysis of lung single-cell sequencing reveals a transitional profibrotic macrophage. Nat. Immunol. 20(2), 163–172 (2019)

21. Carithers, L.J., Ardlie, K., Barcus, M., Branton, P.A., Britton, A., Buia, S.A., Compton, C.C., DeLuca, D.S., Peter-Demchok, J., Gelfand, E.T., Guan, P., Korzeniewski, G.E., Lockhart, N.C., Rabiner, C.A., Rao, A.K., Robinson, K.L., Roche, N.V., Sawyer, S.J., Segre, A.V., Shive, C.E., Smith, A.M., Sobin, L.H., Undale, A.H., Valentino, K.M., Vaught, J., Young, T.R., Moore, H.M., GTEx Consortium: A novel approach to High-Quality postmortem tissue procurement: The GTEx project. Biopreserv. Biobank. 13(5), 311–319 (2015)

22. 1000 Genomes Project Consortium, Auton, A., Brooks, L.D., Durbin, R.M., Garrison, E.P., Kang, H.M., Korbel, J.O., Marchini, J.L., McCarthy, S., McVean, G.A., Abecasis, G.R.: A global reference for human genetic variation. Nature 526(7571), 68–74 (2015)

23. Su, Z., Marchini, J., Donnelly, P.: HAPGEN2: simulation of multiple disease SNPs. Bioinformatics 27(16), 2304–2305 (2011)

24. Huang, Q.Q., Ritchie, S.C., Brozynska, M., Inouye, M.: Power, false discovery rate and winner’s curse in eQTL studies. Nucleic Acids Res. 46(22), 133 (2018)

25. McCarthy, D.J., Campbell, K.R., Lun, A.T.L., Wills, Q.F.: Scater: pre-processing, quality control, normalization and visualization of single-cell RNA-seq data in R. Bioinformatics 33(8), 1179–1186 (2017)

26. Maechler, M., Rousseeuw, P., Struyf, A., Hubert, M., Hornik, K.: Cluster: Cluster Analysis Basics and Extensions. (2021). R package version 2.1.2 — For new features, see the ‘Changelog’ file (in the package source). https://CRAN.R-project.org/package=cluster

27. Benjamini, Y., Hochberg, Y.: Controlling the false discovery rate: A practical and powerful approach to multiple testing. J. R. Stat. Soc. Series B Stat. Methodol. 57(1), 289–300 (1995)

28. Lun, A.T.L., McCarthy, D.J., Marioni, J.C.: A step-by-step workflow for low-level analysis of single-cell RNA-seq data with bioconductor. F1000Res. 5, 2122 (2016)

29. Casale, F.P., Rakitsch, B., Lippert, C., Stegle, O.: Efficient set tests for the genetic analysis of correlated traits. Nat. Methods 12(8), 755–758 (2015)

30. Purcell, S., Neale, B., Todd-Brown, K., Thomas, L., Ferreira, M.A.R., Bender, D., Maller, J., Sklar, P., de Bakker, P.I.W., Daly, M.J., Sham, P.C.: PLINK: a tool set for whole-genome association and population-based linkage analyses. Am. J. Hum. Genet. 81(3), 559–575 (2007)

