## Supplemental Materials for "splatPop: simulating population scale single-cell RNA sequencing data"

### Table of contents

#### Contents

**Table of contents** 1

**Supplementary Tables** 2

**Supplementary Figures** 5

### Supplementary Tables

**Table S1.** Input parameters for the splatPop simulation model

| ID | Estimated | Category | Symbol | Description | Default Value |
|---|---|---|---|---|---|
| **Population parameters** | | | | | |
| pop.mean.shape | yes | population | $\alpha\_m$ | Gamma shape for population-wide gene means | 0.34 |
| pop.mean.rate | yes | population | $\beta\_m$ | Gamma rate for population-wide gene means | 0.008 |
| pop.cv.param | yes | population | $\alpha\_v$, $\beta\_v$ | Data frame containing gamma shape and rate for population-wide gene variance by bin | varies |
| **eQTL parameters** | | | | | |
| eqtl.ES.shape | yes | eQTL | $\alpha\_e$ | eQTL & Gamma shape for eQTL effect sizes | 3.6 |
| eqtl.ES.rate | yes | eQTL | $\beta\_e$ | eQTL & Gamma rate for eQTL effect sizes. | 12 |
| **Single-cell parameters** (estimated with original splatEstimate function) | | | | | |
| mean.shape | yes | single-cell | $\alpha\_sc$ | Shape parameter for the mean gene expression gamma distribution | 0.6 |
| mean.rate | yes | single-cell | $\beta\_sc$ | Rate parameter for the mean gene expression gamma distribution | 0.3 |
| lib.loc | yes | single-cell | $\mu\_L$ | Location parameter for the library size log-normal distribution | 11 |
| lib.scale | yes | single-cell | $\sigma\_L$ | Scale parameter for the library size log-normal distribution | 0.2 |
| out.prob | yes | single-cell | $\pi\_O$ | Probability that a gene is an expression outlier | 0.05 |
| out.facLoc | yes | single-cell | $\mu\_O$ | Location parameter for the expression outlier factor log-normal distribution | 4 |
| out.facScale | yes | single-cell | $\sigma\_O$ | Scale parameter for the expression outlier factor log-normal distribution | 0.5 |
| bcv.common | yes | single-cell | $\varphi$ | Common BCV dispersion across all genes in single-cell data | 0.1 |
| bcv.df | yes | single-cell | $df\_0$ | Degrees of freedom for the BCV inverse chi-squared distribution | 60 |

| | | | | | |
|---|---|---|---|---|---|
| dropout.mid | yes | single-cell | x_0 | Midpoint for the dropout logistic function | 0 |
| dropout.shape | yes | single-cell | k | Shape of the dropout logistic function | -1 |
| **Manual parameters** | | | | | |
| similarity.scale | no | population | s_s | Scaling factor for pop.cv.param.rate, where values larger than 1 increase the similarity between individuals in the population and values less than one make the individuals less similar. | 1 |
| pop.cv.bins | no | population | | Number of gene mean bins to use to estimate CV params | 10 |
| pop.quant.norm | no | population | | T/F if simulated gene means per individual should be quantile normalized to fit the distribution of the single-cell gene mean distribution | TRUE |
| eqtl.n | no | eQTL | | Number (if >1) or proportion of genes to simulate as eGenes | 1 |
| eqtl.dist | no | eQTL | | Maximum distance from center of eGene to eSNP | 1 Mb |
| eqtl.maf.min | no | eQTL | | Minimum minor allele frequency of eSNP | 0.05 |
| eqtl.maf.max | no | eQTL | | Maximum minor allele frequency of eSNP | 0.5 |
| eqtl.group.specific | no | eQTL | | Proportion of eQTL to set as group specific if nGroups >1 | 0.2 |
| eqtl.condition.specific | no | eQTL | | Proportion of eQTL to set as condition specific if nConditions >1 | 0.2 |
| nCells.sample | no | single-cell | | T/F if nCells should be sampled from a gamma distribution for each batch/donor. | FALSE |
| nCells.shape | no | single-cell | | Shape parameter for the nCells per batch per donor distribution. | 1.5 |
| nCells.rate | no | single-cell | | Rate parameter for the nCells per batch per donor distribution. | 0.015 |
| batch.size | no | batch effects | | The number of donors in each pool/batch. | 10 |
| de.prob | no | group effects | π_de | Probability that a gene is DE in a cell group. | 0.1 |

| | | | | | |
|---|---|---|---|---|---|
| de.downProb | no | group effects | | Probability that a group-DE gene is down-regulated. | 0.5 |
| de.facLoc | no | group effects | μ_de | Location (meanlog) parameter for the group-DE factor log-normal distribution. | 0.1 |
| de.facScale | no | group effects | σ_de | Scale (sdlog) parameter for the group-DE factor log-normal distribution. | 0.4 |
| nConditions | no | conditional effects | | The number of conditions/treatments to divide samples into. | 1 |
| condition.prob | no | conditional effects | | Probability that a sample belongs to each condition/treatment group. | c(0.5, 0.5) |
| cde.prob | no | conditional effects | π_cde | Probability that a gene is DE in a conditional cohort. | 0.1 |
| cde.downProb | no | conditional effects | | Probability that a conditional-DE gene is down-regulated. | 0.5 |
| cde.facLoc | no | conditional effects | μ_cde | Location (meanlog) parameter for the conditional-DE factor log-normal distribution. | 0.1 |
| cde.facScale | no | conditional effects | σ_cde | Scale (sdlog) parameter for the conditional-DE factor log-normal distribution. | 0.4 |

**Table S2.** Time (minutes) to simulate populations with N individuals (rows) and N genes (columns), with 100 cells from a single cell-group simulated per individual.

| | | **# genes** | | | |
|---|---|---|---|---|---|
| | | **10** | **100** | **500** | **1000** |
| | **10** | 3.9 | 6.8 | 24.5 | 45.3 |
| | **100** | 24.0 | 28.1 | 47.0 | 69.3 |
| | **500** | 118.5 | 123.8 | 147.5 | 178.5 |
| **# individuals** | **1000** | 236.2 | 244.6 | 273.5 | 312.4 |

### Supplementary Figures

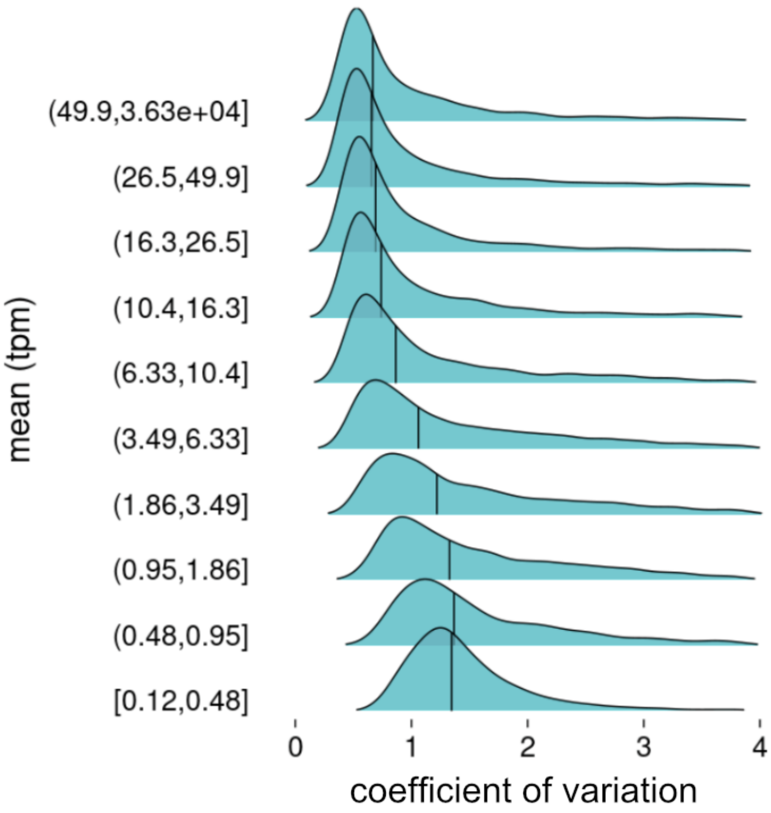


**Figure S1. The gene mean-variance trend.** The distribution of gene expression coefficient of variation between individuals, binned by gene mean across individuals (y-axis). Expression data from GTEx thyroid tissue. The bar indicates the median variance per expression bin. tpm: transcripts per million.

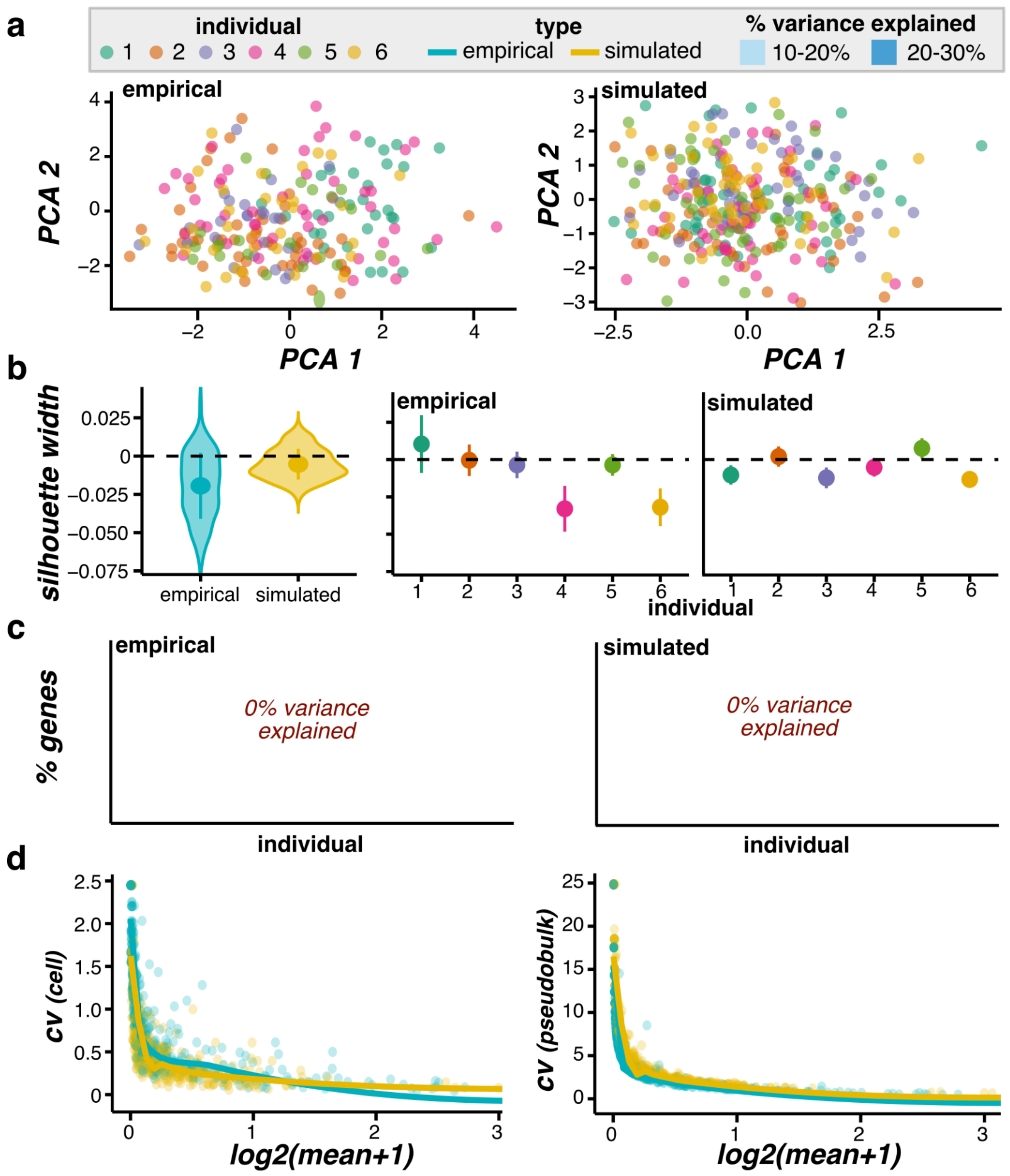


**Figure S2. Simulated compared to empirical 10x-Neuro single-cell RNA-seq data. (a)** PCA plots of cells colored by individual. **(b)** The distribution of cell silhouette widths using the individual as the cluster. The distributions are shown for cells grouped by type (left) and by type and individual (right), with the point and whisker showing the mean and standard deviation. **(c)** The percent of genes (y-axis) with a given percentage of variance explained by individual. Note, no variance in gene expression was explained by individual in the 10x-Neuro empirical or simulated data. **(d)** The mean-variance relationship, with the average gene mean (x-axis) and the gene coefficient of variation (cv; y-axis) calculated across (left) all cells or (right) pseudo-bulk data aggregated by individual. Empirical and simulated: n = 50 cells per individual.

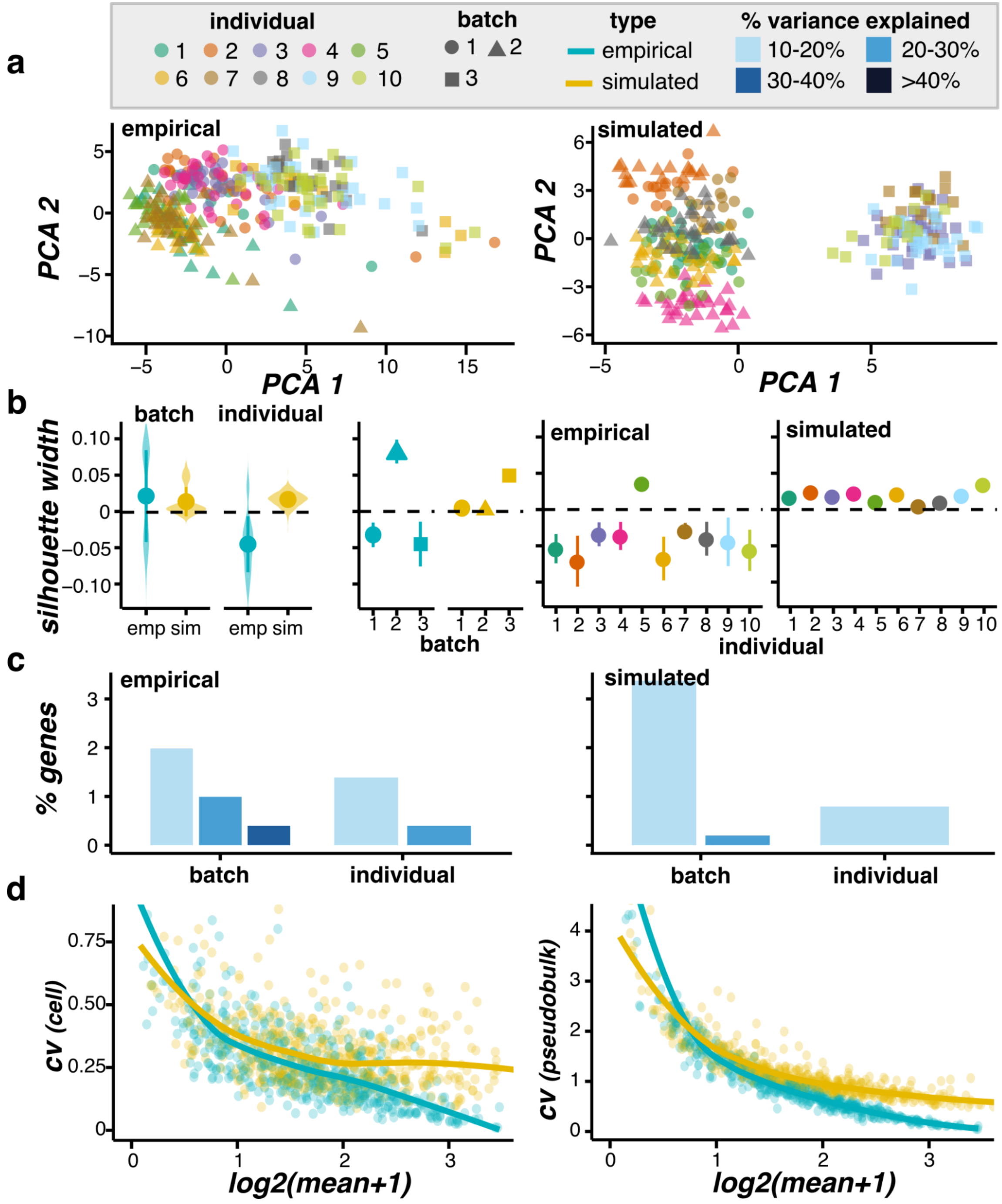


**Figure S3. Simulated compared to empirical SmartSeq2 iPSC single-cell RNA-seq data (ss2-iPSC) from three batches. (a)** PCA plots of cells colored by individual and shaped by batch (max 50 cells shown per individual). **(b)** The distribution of cell silhouette widths using the batch or individual as the cluster. The distributions are shown for cells grouped by type (left) and by type, batch (middle), and individual (right), with the point and whisker showing the mean and standard deviation. **(c)** The percent of genes (y-axis) with a given percentage of variance explained by batch and individual. **(d)** The mean-variance relationship, with the average gene mean (x-axis) and the gene coefficient of variation (cv; y-axis) calculated across (left) all cells or (right) pseudo-bulk data aggregated by individual. Empirical: n= average 52 cells per individual; Simulated: n = 52 cells per individual.

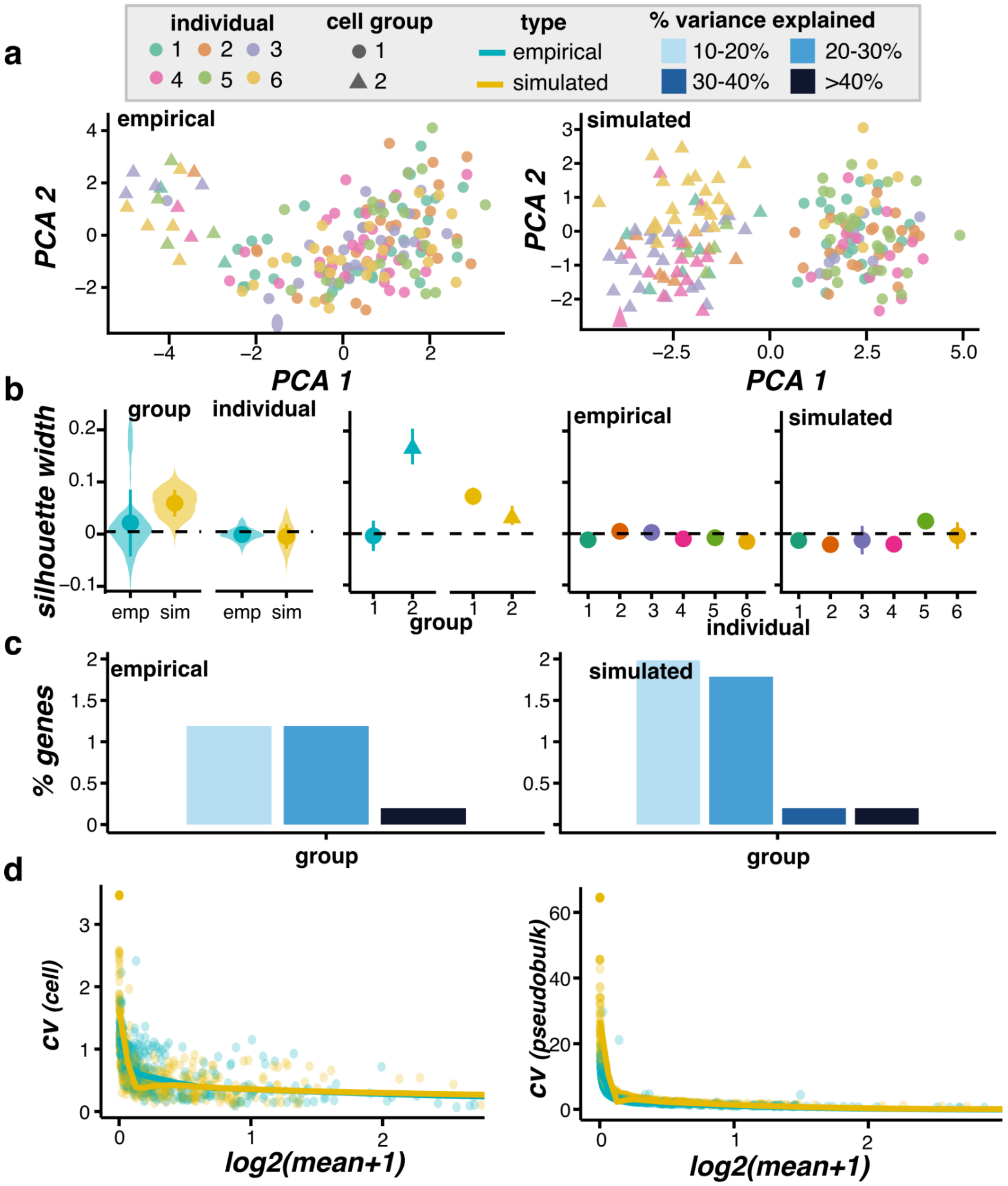


**Figure S4. Simulated compared to empirical 10x-Neuro data from floor plate progenitor and dopaminergic neuron cells. (a)** PCA plots of cells colored by individual and shaped by cell group (max 50 cells shown per individual). **(b)** The distribution of cell silhouette widths using the cell group or individual as the cluster. The distributions are shown for cells grouped by type (left) and by type, cell group (middle), and individual (right), with the point and whisker showing the mean and standard deviation. **(c)** The percent of genes (y-axis) with a given percentage of variance explained by cell group and individual. Note, no variance in gene expression was explained by individual in the 10x-Neuro empirical or simulated data. **(d)** The mean-variance relationship, with the average gene mean (x-axis) and the gene coefficient of variation (cv; y-axis) calculated across (left) all cells or (right) pseudo-bulk data aggregated by individual. Empirical and simulated: n = 50 cells per individual.

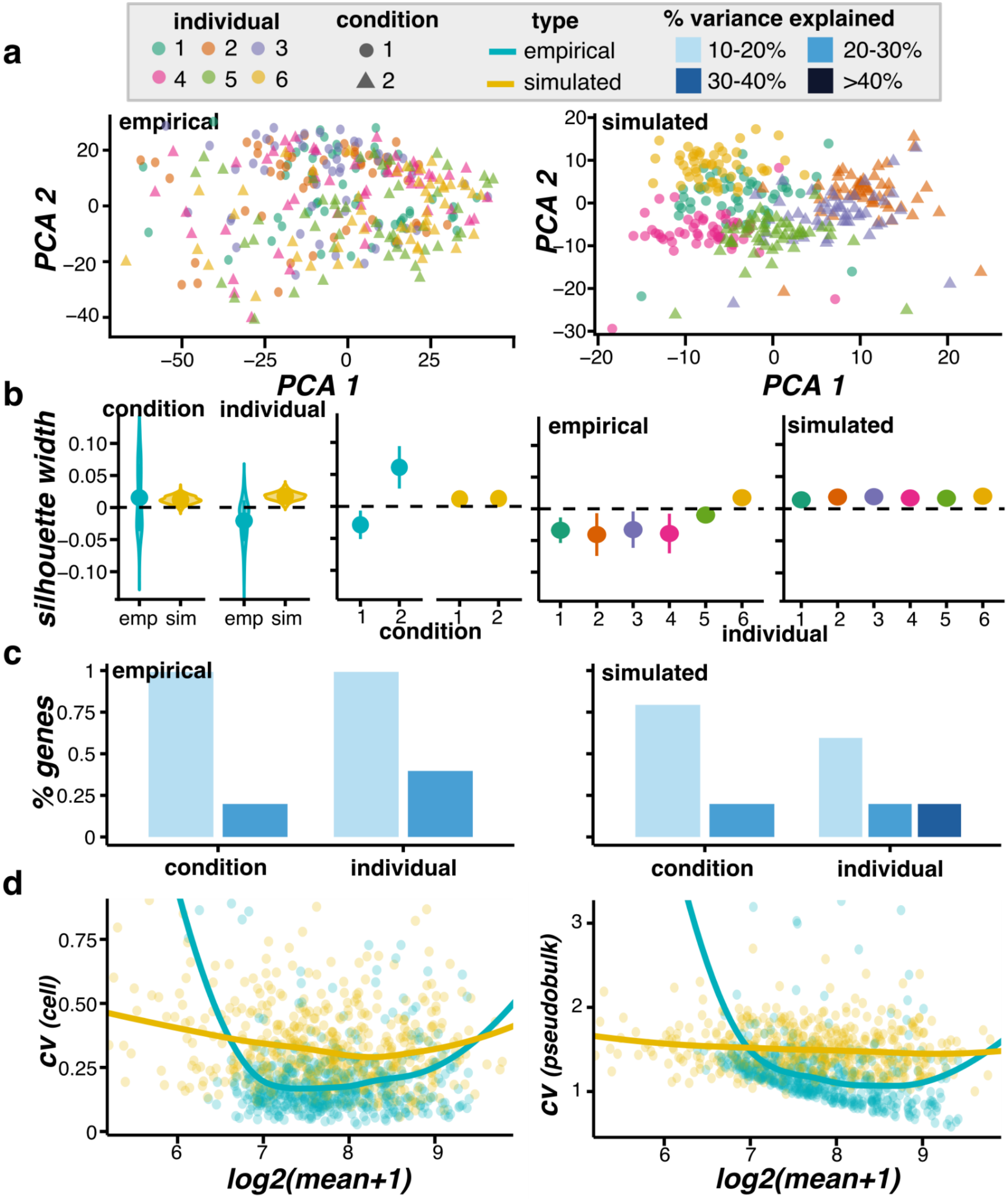


**Figure S5. Simulated compared to empirical 10x-IPF single-cell RNA-seq data from fibroblast cells for 3 healthy and 3 stimulated IPF samples. (a)** PCA plots of cells colored by individual and shaped by conditional group. **(b)** The distribution of cell silhouette widths using the conditional group or individual as the cluster. The distributions are shown for cells grouped by type (left) and by type, conditional group (middle), and individual (right), with the point and whisker showing the mean and standard deviation. **(c)** The percent of genes (y-axis) with a given percentage of variance explained by conditional group and individual. **(d)** The mean-variance relationship, with the average gene mean (x-axis) and the gene coefficient of variation (y-axis) calculated across (left) all cells or (right) pseudo-bulk data aggregated by individual. Empirical and simulated: n = 50 cells per individual.
